# On the role of tail in stability and energetic cost of bird flapping flight

**DOI:** 10.1101/2022.07.12.499802

**Authors:** Gianmarco Ducci, Gennaro Vitucci, Philippe Chatelain, Renaud Ronsse

## Abstract

Migratory birds travel over impressively long distances. Consequently, they have to adopt flight regimes being both efficient - in order to spare their metabolic resources - and robust to perturbations.

This paper investigates the relationship between both aspects, i.e. mechanical performance and stability in flapping flight of migratory birds. Relying on a poly-articulated wing morphing model and a tail-like surface, several families of steady flight regime have been identified and analyzed. These families differ by their wing kinematics and tail opening. A systematic parametric search analysis has been carried out, in order to evaluate power consumption and cost of transport. A framework tailored for assessing limit cycles, namely Floquet theory, is used to numerically study flight stability.

Our results show that under certain conditions, an inherent passive stability of steady and level flight can be achieved. In particular, we find that progressively opening the tail leads to passively stable flight regimes. Within these passively stable regimes, the tail can produce either upward or downward lift. However, these configurations entail an increase of cost of transport at high velocities penalizing fast forward flight regimes.

Our model-based predictions suggest that long range flights require a furled tail configuration, as confirmed by field observations, and consequently need to rely on alternative mechanisms to stabilize the flight.

## I. Introduction

Biological fliers are a source of inspiration for the scientific community. Their capacity to travel over long distances during migrations, their responsiveness to environmental perturbations, and their maneuvering skills are intriguing and inspiring biologists and engineering advances. A particularly outstanding capacity is how they can robustly react to gusts and other perturbations [1, 2]. A foundational study by Smith [3] developed a theory of *evolution of instability*, establishing how inherently unstable flight regimes might have provided a selective advantage for fliers through evolution. Indeed, passively unstable systems are more responsive to changes in command, and this might have facilitated maneuverability for birds. This had to come in parallel with the development of sensory-driven neural circuitries to actively control the flight in order to display stable closed-loop behavior.

Over the last couple of decades, several studies have investigated how such stability might be achieved, with a specific focus on the gliding regime. Thomas and Taylor [4] studied gliding flight and showed that birds use a combination of passive stability — alleviating perturbations without active control — governed by their morphology, and active stabilization from neural pathways to regulate their flight. For example, in gliding gulls, static longitudinal stability is achievable by modulating the opening of the elbow joint over a large range [5, 6, 7]. Cheney et al. [1] investigated the role of wing compliance and tail actuation in order to alleviate perturbations. Ajanic et al. [8] conducted a dedicated study on wing morphing and the mechanism of wing sweep on a propelled gliding robot. For each morphological configuration, the authors estimated the required power to fly. They showed that sweeping the wing backwards and increasing the tail surface was beneficial for longitudinal passive stability, although at the cost of increasing parasitic drag and thus decreasing energetic performance.

By definition, gliding is a flight regime that does not produce the thrust needed to maintain flight altitude over long distances. This is instead possible with the flapping regime. However, studying passive stability of flapping flight requires a dedicated framework to handle the periodic nature of this locomotion regime, and thus the existence of limit cycles instead of fixed equilibrium points. In particular, such a framework must capture the periodic kinematic-dependent nature of aerodynamics forces. Taylor and Thomas [9] pioneered the developments of such a mathematical framework for studying longitudinal stability in flapping flight. They unveiled that flight stability is affected by the location where the mean aerodynamic force is applied with respect to the body center of mass. Taylor and Żbikowski [10] re-defined stability of flapping flight as the asymptotic orbital stability of a limit cycle in the phase space. Following this approach, Dietl and Garcia [11] used for the first time Floquet theory [12] to study the phase space limit cycle described by the equations of motion of an an artificial flying device, namely an ornithopter. We recently leveraged the same formalism to characterize the passive stability of large migratory birds [13]. Our framework builds upon a morphing quasi-steady lifting line model to compute the aerodynamic forces, and a multiple-shooting algorithm to identify limit cycles. The aerodynamic effects induced by the tail were not captured within this previous work, and the flight was consequently characterized by an unstable mode in pitch whatever the wing configuration.

Moreover, in flapping flight, the wing kinematics has an important impact on power consumption. In an analysis on pigeons, Parslew [14] suggested that particular kinematics modes might be selected in specific flight regimes for energy saving purposes. Colognesi et al. [15] also showed a dependency between power requirement and key parameters of the wing kinematics, specifically the wingbeat amplitude.

Although the role of the tail has been studied in gliding regimes, and the influence of wing kinematics has been studied to assess the performance in flapping regimes, no study to date combined both in a whole body characterization of flapping gaits. The objective of this paper is to provide such a complete modeling of flapping flight that can simultaneously assess the influence of wing kinematics and tail morphology in stability and energetic performance. Such a framework would be of direct relevance to challenge biological hypotheses suggesting an evolution towards passively unstable flight regimes for enhancing maneuverability. Based on observations with real birds [3], we hypothesize that opening the tail should inherently lead to passively stable flight regimes, at the price of an increased energetic cost.

The rest of the paper is structured as follows: Section II develops the model capturing the bird biomechanics and its aerodynamic interactions with the environment. Section III overviews the multiple-shooting algorithm used to find the steady conditions and evaluate flight stability. It also shows how the mechanical power of flight is calculated and reports the numerical parameters used in the experiments. Finally, Section IV reports the results of the numerical investigations, and Section V concludes the paper.

## II. Dynamical model of bird flight

In this section, we introduce flapping flight dynamics and describe the bird model used in our computational framework. In particular we introduce the equations of longitudinal dynamics and describe the wing model and its kinematics. A particular emphasis is put on capturing the tail morphing.

### i. Equations of motion

Flight dynamics is restricted to the longitudinal plane and thus the bird main body is captured as a rigid-body with three degrees of freedom, i.e. two in translation and one in rotation. This model preserves symmetry with respect to this plane, without any lateral force and moment. The aerodynamic model of the wing relies on the theory of quasi-steady lifting-line [16]. Additionally, the present work does not account for the so-called inertial power, required for accelerating and decelerating the wing over a flapping period, since wing inertia is neglected [10, 11, 17]. This inertial power was shown to be negligible in fast forward flight conditions, in comparison to the other contributions to actuation power [18].

The body is thus modeled with a mass *m*_*b*_ and a rotational inertia *I*_*yy*_ about its center of mass. The equations of motion are expressed in the body frame *G*(*x*′, *y*′, *z*′) with unit vectors (**ê**_*x*′_, **ê**_*y*′_, **ê**_*z*′_), and an origin located at the center of mass, as pictured in Figure 1. The state space vector is thus

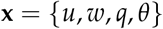

where *u* and *w* are the body velocities along the *x*′− and *z*′−axis and *θ* and *q* are the pitch angle and its time derivative about the *y*′−axis, respectively. Consequently, the equations of motion read [11, 13, 19]

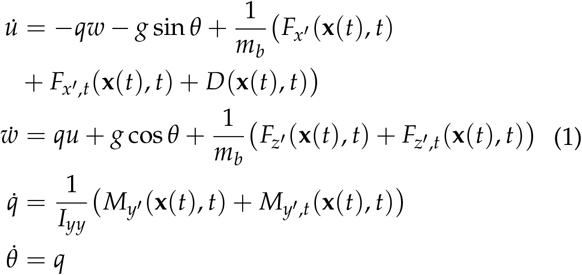

**Figure 1:**
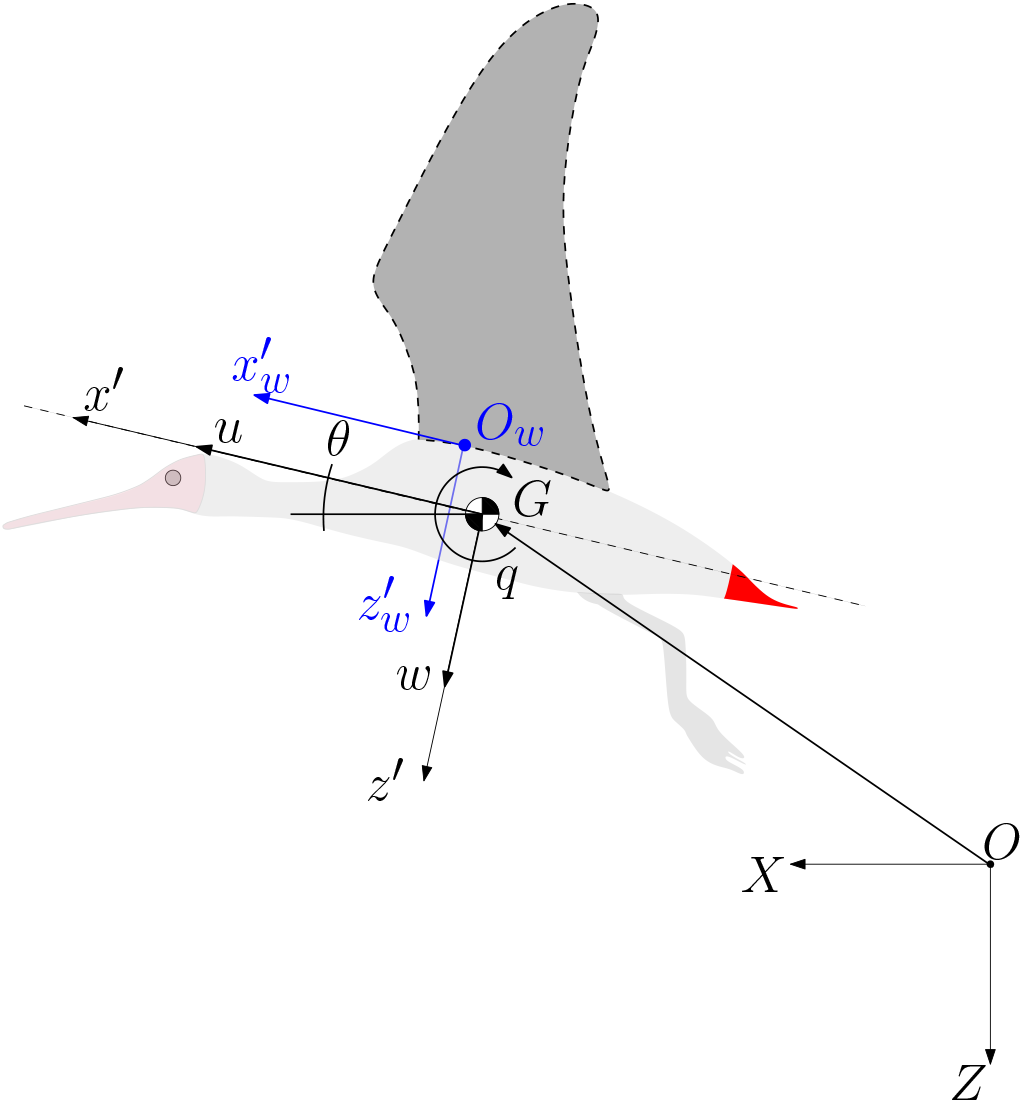
Bird model for describing the flight dynamics in the longitudinal plane. The state variables are expressed with respect to the moving body-frame located at the flier’s center of mass G(x′, z′). These state variables are the component of forward flight velocity, u, the velocity component of local vertical velocity, w, the orientation of this body-centered moving frame with respect to the fixed frame, θ and its angular velocity, q. A second frame 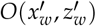 is used to compute the position of the wing, relative to the body. The wings (dark gray) and the tail (red) are the surfaces of application of aerodynamic forces.

The forcing terms in Eq. 1 are the aerodynamic forces and moments applied to the wing (namely *F*_*x*′_, *F*_*z*′_, and *M*_*y*′_) and to the tail (*F*_*x*′*t*_, *F*_*z*′*t*_, and *M*_*y*′, *t*_). The whole drag is captured by an extra force *D* that sums contributions due to the body *D*_*b*_, the skin friction of the wing (wing profile) *D*_*p,w*_, and the skin friction of the tail (tail profile) *D*_*p,t*_. These terms are described in detail in the next sections.

### ii. Wing model

The bird has two wings. Each wing is a rigid poly-articulated body, comprising the bird arm, forearm and hand, as pictured in Figure 2. Each segment is actuated by a joint to induce wing morphing. We refer to [13, 15] for a complete description of this wing kinematic model.

**Figure 2:**
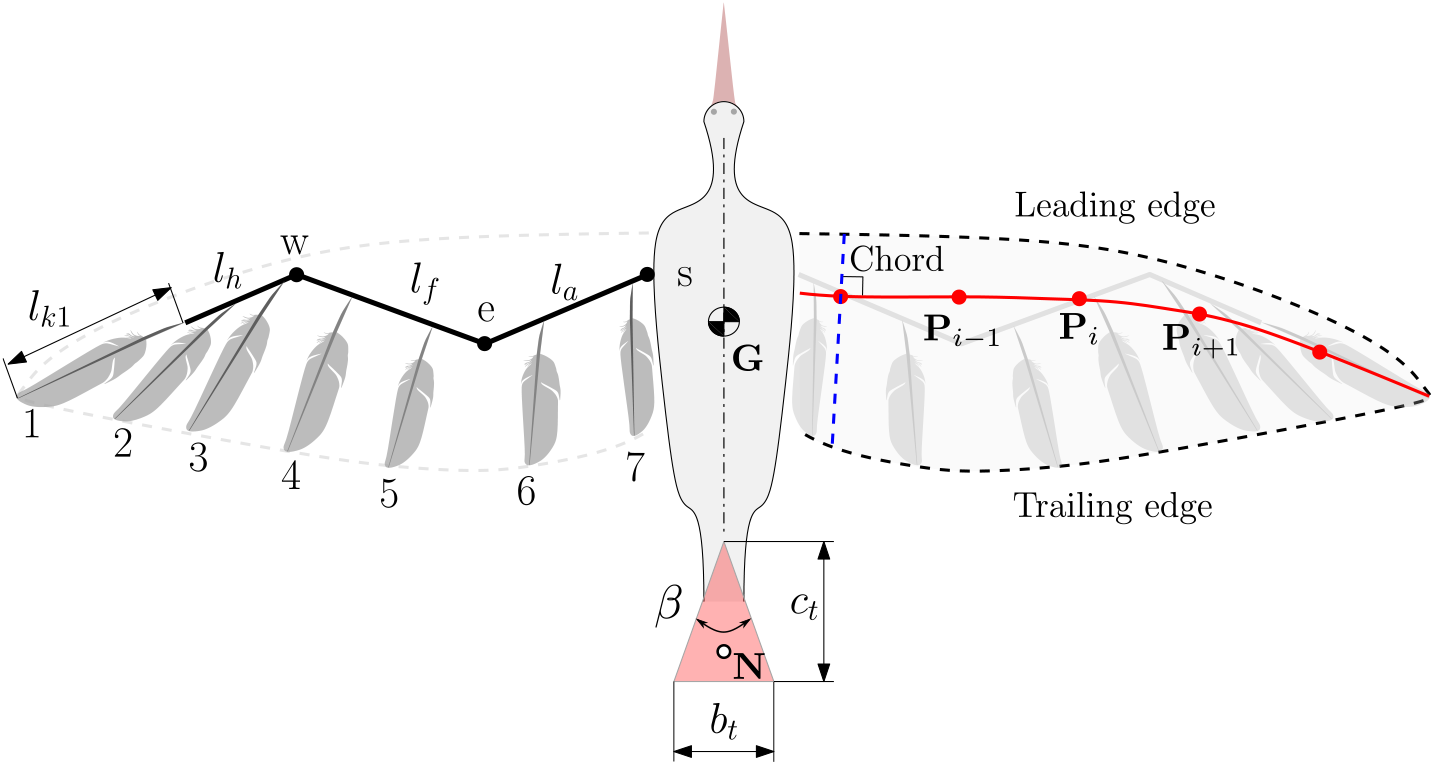
Top view of the bird model. The left wing emphasizes a cartoon model of the skeleton. The shoulder joint **s** connects the wing to the body via three rotational degrees of freedom (RDoF), the elbow joint **e** connects the arm with the forearm via one RDoF and the wrist joint **w** connects the forearm to the hand via two RDoF. Each feather is attached to a bone via two additional RDoF, except the most distal one “1” which is rigidly aligned with the hand. The right wing further emphasizes the lifting line (red) which is computed as a function of the wing morphing. The aerodynamic forces generated on the wing are computed on the discretized elements P_i_. The tail is modeled as a triangular shape with fixed chord c_t_ and maximum width b_t_ that can be morphed as a function of its opening angle β.

Each joint is kinematically driven to follow a sinusoidal trajectory specified as:

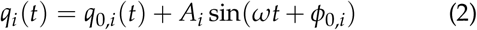

with *ω* = 2*π f* and *f* being the flapping frequency which is identical for each joint, *q*_0,*i*_ being the mean angle over a period (or offset), *A*_*i*_ the amplitude, and *φ*_0,*I*_ the relative phase of joint *i*. A complete wingbeat cycle is therefore described through a set of 19 kinematic parameters, including the frequency *f*.

We assume that the wing trajectory is rigidly constrained, and therefore we do not need to explicitly solve the wing dynamics. Under this assumption, the motion generation does not require the computation of joint torques. The model further embeds seven feathers of length *l*_*ki*_ in each wing. They are attached to their respective wing bones via two rotational degrees of freedom allowing them to pitch and spread in the span-wise direction. These two degrees of freedom are again kinematically driven by relationships that depend on the angle between the wing segments [13]. This makes the feathers spreading and folding smoothly through the wingbeat cycle. In sum, the kinematic model of the wing yields the position of its bones and feathers at every time step. This provides a certain wing morph-ing from which the wing envelope (leading edge and trailing edge) can be computed (see Figure 2). From the wing envelope, the aerodynamic chord and the lifting line are computed. The lifting line is the line passing through the quarter of chord, which is itself defined as the segment connecting the leading edge to the trailing edge and orthogonal to the lifting line (Figure 2). This extraction algorithm is explained in detail in [15]. In order to calculate the aerodynamic forces, the angle of attack of the wing profile has to be evaluated. Each wing element defines a plane containing the lifting line and the aerodynamic chord as pictured in Figure 3. The orientation of the plane is identified by the or-thogonal unit vectors (**ê**_*n*_, **ê**_*t*_, **ê**_*b*_), where **ê**_*n*_ is the vector perpendicular to the plane and **ê**_*t*_ is the tangent to the lifting line. To compute the effective angle of attack, the velocity perceived by the wing profile is computed as the sum of the velocities due to the body and wing motion, and the velocity induced by the wake. The first contribution, **U**, accounts for

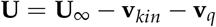

where **U**_∞_ = *u***ê**_*x*′_ + *w***ê**_*z*′_ is the actual flight velocity, **v**_*kin*_ is the relative velocity of the wing due to its motion, and **v**_*q*_ is the component induced by the angular velocity of the body *q* and calculated as

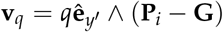

**Figure 3:**
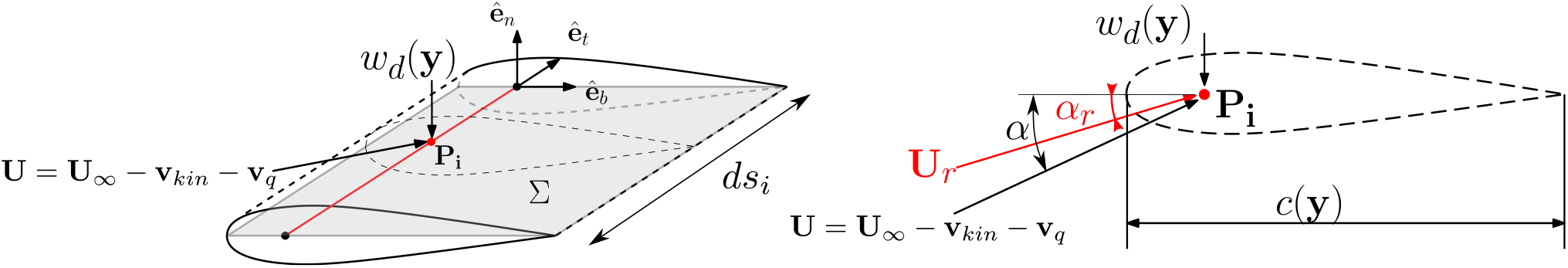
Left:Wing element i between two wing profiles, identifying a plane Σ containing the lifting line (red). Right: Cross section of the wing element, containing the chord point **P**_**i**_ where the velocities are computed.

This velocity vector **U** defines the angle *α*, as pictured in Figure 3.

The second contribution is due to the induced velocity field by the wake, i.e. the downwash velocity *w*_*d*_, and acting along the normal unit vector −*w*_*d*_**ê**_*n*_. The resulting effective angle of attack, *α*_*r*_, is thus

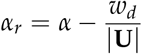

The downwash velocity *w*_*d*_ is computed according to the Biot-Savart law [16], assuming the wake being shed backward in the form of straight and infinitely long vortex filaments at each time step of the simulation [13, 15]. Once the downwash is evaluated, it is possible to evaluate the circulation, and consequently the aerodynamic force and moment acting at the el-ement *P*_*i*_, i.e. *F*_*x*′,*i*_ (**x**(*t*), *t*), *F*_*z*′,*i*_ (**x**(*t*), *t*), *M*_*y*′,*i*_ (**x**(*t*), *t*), as explained in detail in [13]. We use the thin airfoil theory for the estimation of the lift coefficient, with a slope of 2*π* that saturates at an effective angle of attack *α*_*r*_ of ±15^°^.

### iii. Drag production by body and wing

The main body and the wings induce drag that should be accounted for in a model aiming at characterizing energetic performance. Body-induced drag is named parasitic because the body itself does not contribute to lift generation, and only induces skin friction and pressure drags around its envelope [20]. The total body drag is:

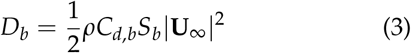

where *ρ* is the air density. We implemented the model described by Maybury [20] to compute the body drag coefficient *C*_*d,b*_. This depends on the morphology of the bird and the Reynolds number *Re* according to

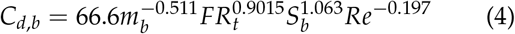

with *S*_*b*_ and *FR*_*t*_ are respectively the frontal area of the body and the fitness ratio of the bird, and both of them can be estimated from other allometric formulas i.e. [20, 21].

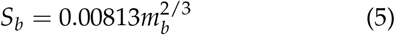

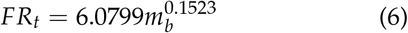

The Reynolds number 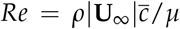 is calculated with the reference length of the mean aerodynamic chord 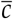, with *µ* being the dynamic viscosity. This model is found to be suitable for Reynolds number in the range 10^4^ − 10^5^ [20]. Another source of drag is the profile drag due to friction between the air and the feathers on the wings. It is the sum of the profile drag at each section along the wingspan, i.e.

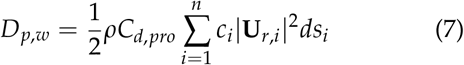

with *c*_*i*_ the chord length, *ds*_*i*_ the length of the lifting line element along the wingspan, and **U**_*r,i*_ the velocity at the wing section *i* accounting for the body velocity, the kinematics velocity of the wing and the downwash velocity (Figure 3). We used a value of profile drag of *C*_*d,pro*_ = 0.02 and this is assumed to be constant over the wingspan and throughout the flapping cycle [22].

### iv. Tail model

Since the wingspan of the tail is of the same magnitude as its aerodynamic chord, here the lifting line approach cannot be used [16]. Therefore, the tail is modeled under the slender delta wing theory, as a triangular planform [23]. Its morphology is illustrated in Figure 2 and defined via the opening angle *β* and the chord *c*_*t*_. This latter parameter is kept constant, thus the tail span is controlled via *β* from the trigonometrical relationship

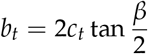

The main limitation of this framework is the low range of angles of attack (*α*_*tail*_ < 5^°^) within which it provides accurate results [24]. This limitation is valid in our context of fast forward flight, where the bird flight is straight, horizontal and the forward velocity *u* is much larger than the vertical one *w*. The velocity component acting on the tail-like surface is

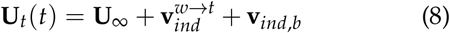

where the term 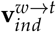is the velocity acting on the tail, induced by vortex filament shed by the wing calculated according to Biot-Savart law [16], and

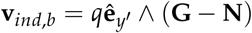

is the velocity induced by the body rotation, with (**G** −**N**) the vector between the center of mass of the body (**G**) and the point of application of the forces on the tail (**N**) taken at two third along the tail chord, as illustrated in Figure 2. The forces generated by the tail are computed according to [23]

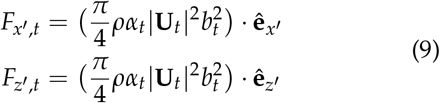

These forces are applied at point **N**. In addition, adding this tail-like surface introduces another source of drag that needs to be accounted for. This parasitic drag contribution is, according to [23]

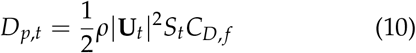

where *S*_*t*_ is the tail planar surface and *C*_*D, f*_ the dimensionless friction coefficient.

## III. Methods

This section reports how the dynamical model described in Section II is used in order to identify steady and leveled flapping flight regimes, study their stability, and assess their energetic performance.

### i. Numerical framework

Due to the fact that Equations 1 are driven by time-varying forces and periodic actuation, they do not converge to steady equilibrium points [11, 25]. Instead, the steady condition corresponds to a limit cycle of this model, and its stability can be assessed via dedicated tools. The numerical identification of such a periodic orbit and the characterization of its stability are challenged by the fact that this orbit is unknown a priori. In previous work, we developed a method to achieve both at the same time via a multiple shooting algorithm. This framework has been released as a Python toolbox called `multiflap` [13]. It takes as input a set of ordinary differential equations (ODE) of the form

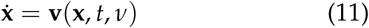

with **x** being the state variables, **v** a vector field describing the system dynamics, *t* the time and *ν* a set of configuration parameters. A periodic orbit is thus a particular solution of this set of ODE such that

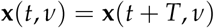

with *T* being the cycle period. Such a periodic solution, defines a steady flight regime.

The stability of such closed orbits can also be assessed through the long term response to a perturbation: if the perturbed trajectory converges back to the orbit, then it is stable, and vice-versa. We perform this stability analysis via the Floquet theory [12]. Stability of the set of equations is governed by the eigenvalues of the so-called Floquet matrix 𝕁, also known as Floquet multipliers, Λ_*i*_. This Floquet matrix maps perturbations within an infinitesimal sphere around a point of the limit cycle (**x**_**0**_, *t*_0_) into an ellipsoid after a time *T* equal to the period of the orbit. Stretching or contracting ratios of the principal axes of this first order transformation are governed by the Floquet Multipliers. Floquet multipliers have the property of being invariant along the limit cycle, whereas the Floquet matrix and its eigenvectors depend on it. Concretely, the Floquet matrix 𝕁 can be calculated as the solution of the variational Equation:

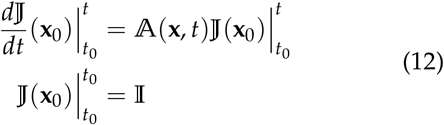

where the matrix

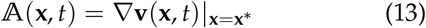

is called the Stability Matrix [26] and is *T*-periodic on the limit cycle. If the absolute values of all Floquet multipliers Λ_*i*_ are smaller than one, the corresponding periodic orbit is stable. Conversely, if the absolute value of at least one multiplier is larger than one, the corresponding orbit is unstable and the perturbation spirals out of the limit cycle along the corresponding eigendirection(s). This framework provides another important feature, namely the stretching/contracting rate per unit of time, or Floquet exponent, *λ*_*i*_ [26, 27]

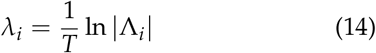

### ii. Application to steady and level flight

In the present study, we leverage the aforementioned multiple-shooting algorithm, for seeking steady flight regimes within the following parametric space

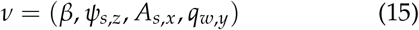

where *β* is the tail opening angle, *ψ*_*s,z*_ is the shoulder sweep offset, *A*_*s,x*_ is the wingbeat amplitude, and *q*_*w,y*_ is the mean rotation angle of the wing profiles of the forearm about the axis *y*, see Figure 4. The other parameters defining the wing kinematics are kept fixed to values similar to those reported in [13].

**Figure 4:**
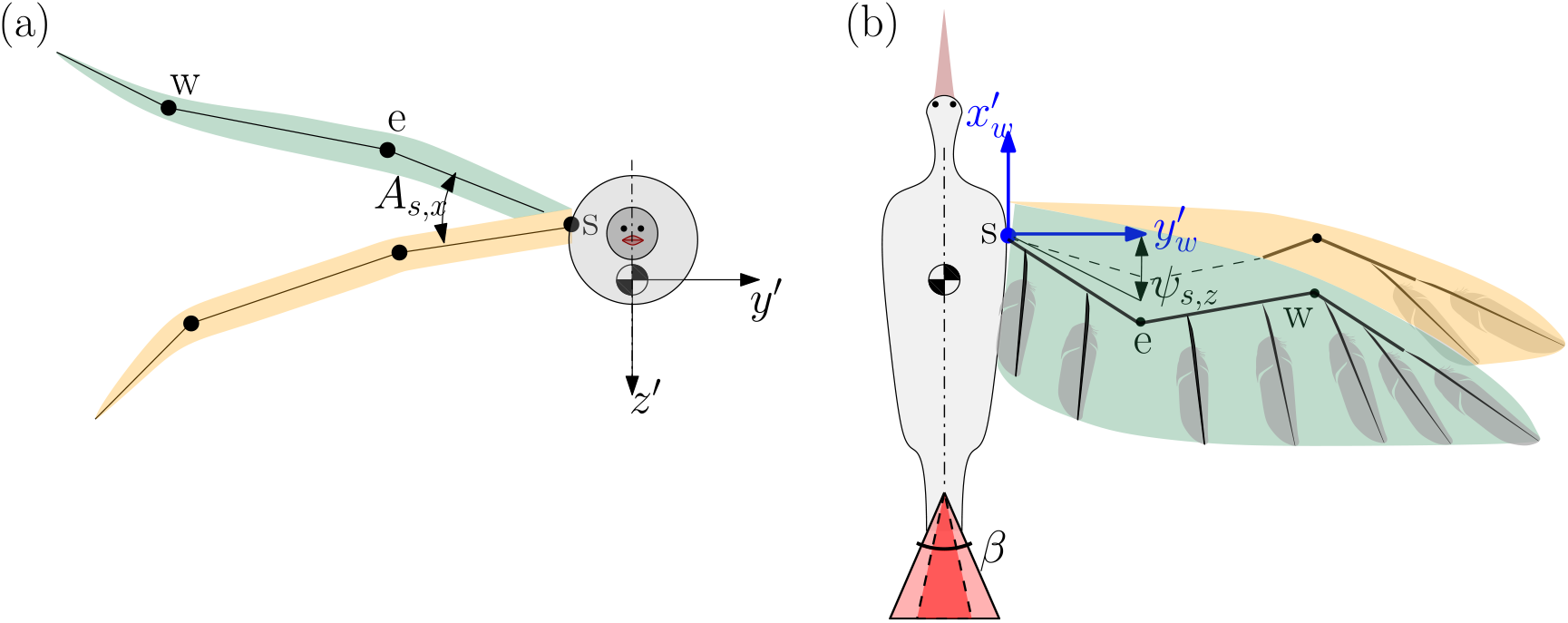
(a): Front view of the bird model. The wingbeat amplitude A_s,x_ about the x′-axis is the main kinematic parameter governing the wing amplitude of movement. (b): Top view of the bird model. The shoulder sweep offset ψ_s,z_ captures the average angle of the arm bone with respect to 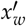 over the period. The angle β captures the magnitude of tail opening.

Previous studies have shown that these four parameters decisively govern the flight regime in bird flapping and gliding modes. The tail opening *β* and shoulder sweep offset *ψ*_*s,z*_ influence flight stability, since these are the parameters having a paramount influence on the generation of pitching moment. Then, the shoulder wingbeat amplitude *A*_*s,x*_ has a direct impact on thrust production and therefore on airspeed and power consumption [28]. The last parameter *q*_*w,y*_ modulates the generation of lift [13, 15].

On top of seeking for steady flight regimes, it is important to identify those corresponding to level flight, i.e. with the bird flying at a constant altitude. This level flight condition thus corresponds to an average mean vertical velocity being equal to zero over the period, in a fixed reference frame. Figure 1 shows that this instantaneous vertical velocity can be computed as

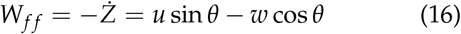

Concretely, satisfying the level flight condition isolates a three-dimensional manifold within the four-dimensional parametric space of Equation 15. Finding this manifold is done by searching the value of the parameter *q*_*w,y*_ that corresponds to level flight for all possible combinations of the three other parameters *β, ψ*_*s,z*_ and *A*_*s,x*_. In other words, we report here only the limit cycles that belong to the manifold satisfying

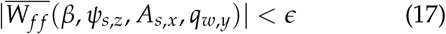

with *ϵ* = 5 · 10^−3^*ms*^−1^, which corresponds to a maximum vertical deviation of 1*mm* per flapping cycle.

### iii. Power Consumption and Cost of Transport

Each limit cycle corresponds to a particular flapping gait with its own mechanical power consumption and the corresponding cost of transport.

Since inertial power for accelerating and decelerating a wing is neglected, the actuation power produced by the wing joints is exactly equal to the power transfered by this wing to the environment, i.e.

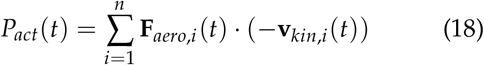

where **v**_*kin,i*_(*t*) is the velocity of the lifting line computed at the discretized point *P*_*i*_ and time *t*, and *F*_*aero,i*_(*t*) is the corresponding aerodynamic load on the wing element *i*, computed by the quasi-steady lifting line model as explained in Section II. The mean power consumption over one flapping period is thus

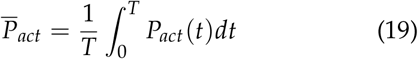

Another important metric to assess locomotion performance is the so-called Cost of Transport (CoT), i.e. a dimensionless ratio equal to the mechanical work produced by the actuators to transport a unit of body weight across a unit of distance [29]. Here, it is thus defined as

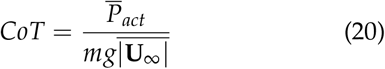

with 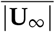 being the magnitude of the flight speed of the corresponding limit cycle, averaged over one period.

### iv. Parametrization of bird morphology and wing kinematics

The above framework and concepts are here embodied in a model of the northen bald ibis (*Geronticus eremita*). The length of bones and feathers are set up accordingly. To the best of our knowledge, the precise wingbeat kinematics of this particular bird have not been reported in the literature. Consequently, we follow the same approach as in [13], consisting in scaling up the detailed kinematic pattern of another bird [30] to the morphology of ours. The morphological and kinematic parameters used to describe the bird are reported in Appendix A.

This study is performed within the following parametric space: tail opening *β* ∈ [0°, 45°], wingbeat amplitude *A*_*s,x*_ ∈ [29°, 45°] and sweep offset *ψ*_*s,z*_ ∈ [9°, 15°]. The parametric space is meshed with an uniform grid spaced along *ψ*_*s,z*_ and *A*_*s,x*_ with a step size of 0.5°, and a step size of 1° along *β*. This resulted in 19,734 possible flight configurations. In the results, we report all solutions satisfying Equation 17, with the addition of two exclusion criteria. First, we excluded limit cycles that do not correspond to fast forward flight. In [31], this corresponds to flying modes 4 and 5 and requires the body pitch angle to stay close to the horizon tail configuration. Concretely, we excluded from the results limit cycles corresponding to a mean body pitch angle larger than 6° in absolute value. Second, we excluded limit cycles corresponding to biologically incompatible kinematics. This was implemented by excluding limit cycles with a mean rotation angle of the wing *q*_*w,y*_ larger than 12° in absolute value. Indeed, remembering that the related amplitude of this joint was fixed to 30° (Table 2), this criteria excluded solutions corresponding to geometrical rotation of the forearm larger than ±42° in absolute value, which we considered to be not physiologically consistent.

## IV. Results

In this section, we report the results of the systematic exploration of the gait parametric space. First, the locus of the solutions is reported, i.e. the set of parametric values for which a limit cycle has been identified. Next, three representative limit cycles are analyzed in detail: the first one with a completely furled tail (*β* = 0), and the both other ones with an open tail (*β* = 40°). Finally, these solutions are assessed in terms of energetic expenditure, quantified by the CoT.

### i. Locus of solutions

Among the 19,734 possible parametric configurations, our algorithm detected 5,604 steady leveled limit cycles. The locus of these identified solutions is pictured in Figure 5.

**Figure 5:**
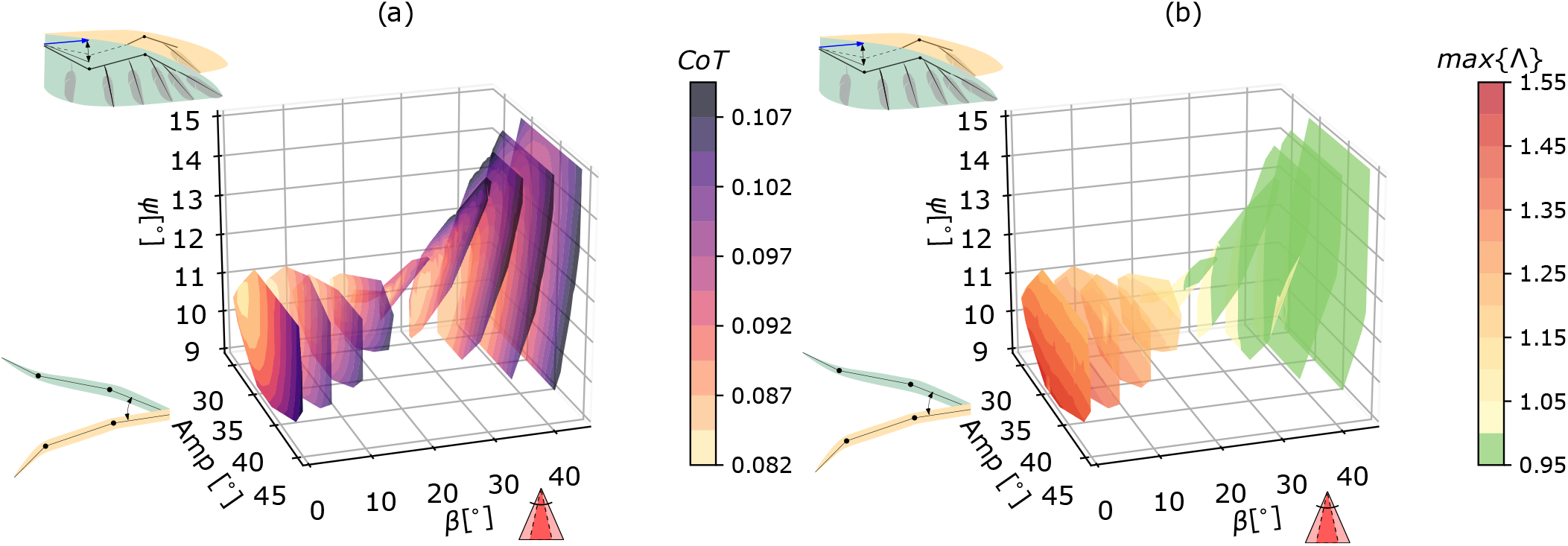
Locus of the steady and leveled solutions in the gait parameter space. Colored surfaces are representing slices of this locus, at every 5^°^ of tail opening. (a): Cost of Transport. (b): Stability indicator captured via the largest Floquet multiplier. Unstable limit cycles are represented with different shades of yellow-to-red, and stables ones are in green; the transition appears around β = 25^°^.

Figure 5(a) shows the CoT at equally-spaced planes of tail opening angle *β*, projected in the parameter space. The CoT progressively increases as the tail spreads out. It further displays a higher gradient with respect to both other parameters for a given opening angle. This reveals that this cost of transport is sensitive to the kinematic parameters governing wing movements. Moreover, with lower tail opening angles, the CoT gradient is lower and less sensitive to changes in kinematics. Figure 5(b) illustrates the stability transition that occurs as a function of the tail opening. The bifurcation point happens for a value around *β* = 25°. The largest Floquet multipliers in the stable region are however never smaller than about 0.96, corresponding to a largest stable Floquet exponent being equal to (see Equation 14)

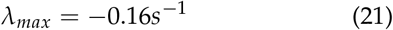

Solving Equation 14 for *t*, the time taken for halving a perturbation is therefore

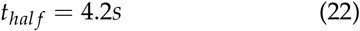

which corresponds to about 17 flapping periods.

As revealed by the quasi-horizontal stripes of uniform colors in Figure 5(b), the shoulder amplitude has a marginal effect on stability — because of its marginal role on the distribution of nose up/down pitching moment — in contrast to the tail opening and sweep angle.

### ii. Comparison between furled and open tail solutions

In this section, three representative limit cycles are further investigated: one corresponding to a tail completely furled (*β* = 0) and the other ones to a tail opening of *β* = 40°. These reference limit cycles are selected to have the same resulting forward flight velocity, i.e. 14*ms*^−1^. The whole set of corresponding parameters is reported in Table 1.

**Table 1:** Parameters for the three representative limit cycles studied in more detail, corresponding to one unstable and two stable flight regimes, respectively, at a forward flight velocity of 14ms^−1^.

|  | (a) | (b) | (c) |
| --- | --- | --- | --- |
| $\beta[^\circ]$ | 0 | 40 | 40 |
| $\psi_{s,z}[^\circ]$ | 10.5 | 14.5 | 13.5 |
| $A_{s,x}[^\circ]$ | 39.5 | 42 | 44 |
| $q_{w,y}[^\circ]$ | 2.3 | -5.5 | -9.67 |

The free-body diagram of these configurations is illustrated in Figure 6-top panel. The actual pitching moment characterizing the limit cycle solutions is reported in Figure 6-middle panel. In case (a) (furled tail), the wing must generate an equal amount of nose-up and nose-down moment to guarantee the existence of a limit cycle. Both open-tail configurations, corresponds to different configurations of momentum equilibrium. In case (b), the wing contributes for nose-down moment (on average) balanced by the nose-up moment (on average) of the tail. Conversely, in the case (c), the wing contributes for nose-up moment (on average) balanced by the nose-down moment (on average) of the tail. All these configurations exhibit a similar limit cycle regarding the phase space trajectory, although the former has one unstable mode in pitch stability gov-erned by a Floquet multiplier equal to Λ = 1.33. The open tail cases are both stable since their largest multiplier has a magnitude equal to about Λ = 0.96. All multipliers are pictured in the inset plot of the middle panel of Figure 6.

**Figure 6:**
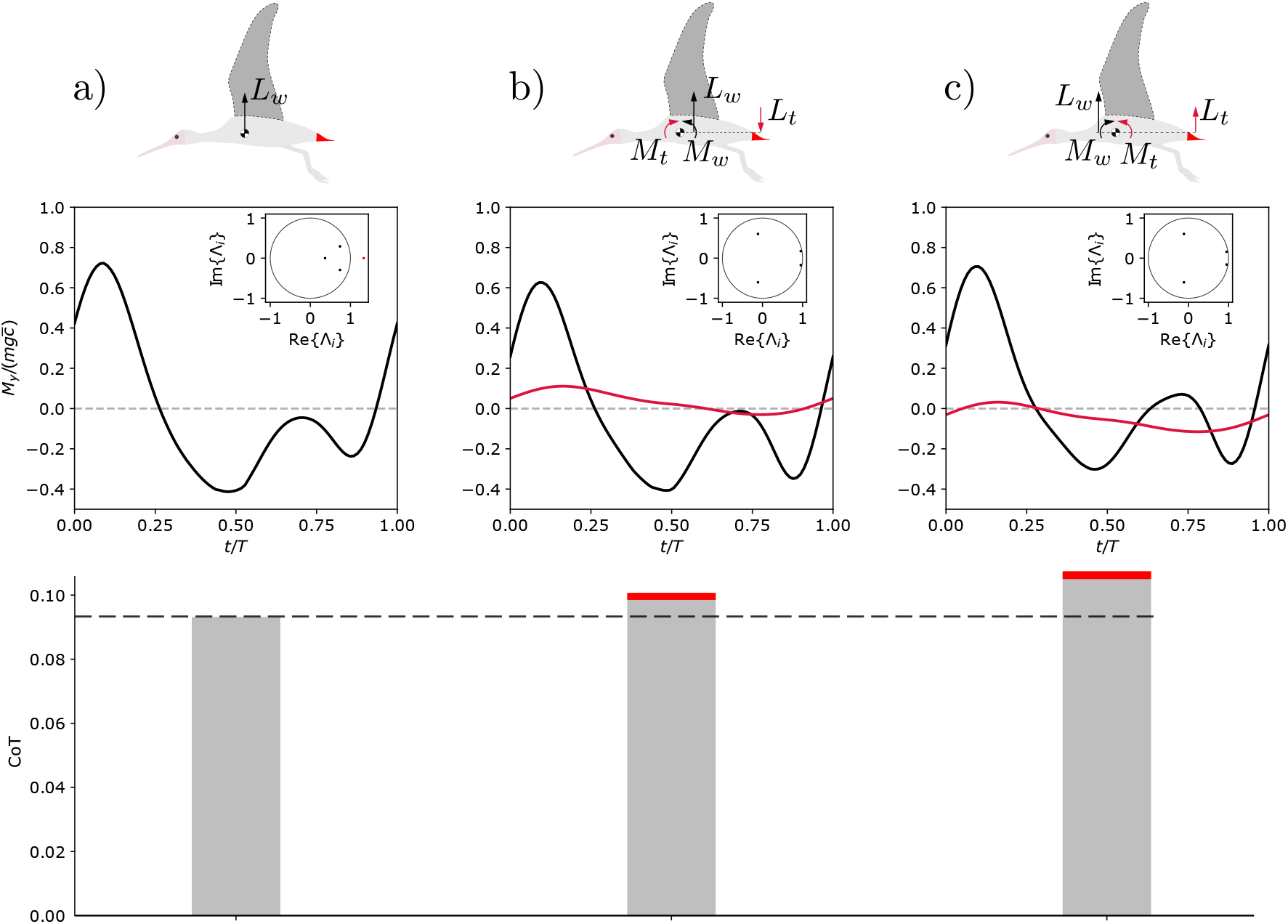
Characterization of three representative limit cycles: one with furled tail (a), and two with a tail opened with an angle β = 40^°^ (b, c). The upper panel represents the free-body diagram of the three different flight configurations. **Case (a):** The pitching moment is only due to the wing movement, and averages at zero. This flight regime is characterized by an unstable mode, highlighted by an eigenvalue larger than 1 (inset middle panel). The bottom panel gives the CoT for this flight configuration. **Case (b):** The average pitching moment M_w_ due to the wing lift (L_w_) is negative (nose down) and the average moment due to the tail lift, M_t_, is positive (nose up). This solution is stable as all the eigenvalues are smaller than 1 in absolute value (inset middle panel). The CoT is quantified in the bottom panel, and with the contribution due to dissipative forces acting on the tail being highlighted in red. **Case (c):** The average pitching moment due to the wing lift (L_w_) is positive (nose up) and the average moment due to the tail lift is negative (nose down). This solution is also stable as all the eigenvalues are smaller than 1 in absolute value (inset middle panel).

The power required to achieve level flight in the furled tail case (a), averaged over one wingbeat cycle, is equal to 15.4*W*. In case (b), it is equal to 16.7*W* with a contribution to to the tail-parasitic drag of about 0.4*W*, while in case (c) it is equal to around 17.9*W* with a power dissipated by drag-induced forces in the tail of about 0.4*W*. This power assessment is pictured adimensionally in the bottom panel of Figure 6, where the red stripes corresponds to the power dissipation from the tail. There is thus a trade-off between robustness to perturbations — characterized by passive stability — and performance — characterized by the required mechanical power.

These three representative limit cycles have been perturbed by an upward gust along the local *z*′-axis. The gust is modeled as a Gaussian signal *w*_*g*_ in the form:

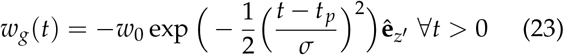

with *t*_*p*_ = 0.25*s* and *σ* = 0.05*s*. The intensity of the gust varies, in order to observe comparable effects in phase space. In the unstable case (a), *w*_0_ = 0.1*ms*^−1^, whereas in the stable cases (b) and (c) *w*_0_ = 1*ms*^−1^. The dynamic response of these three configurations is captured by the black solid lines in Figure 7. Figure 7(a) shows a quick separation from the limit cycle condition (red curves), driven by the unstable Floquet multiplier. Figure 7(b) and (c) show a passively stable response to the perturbation as all the Floquet multipliers are smaller than 1 in both cases. This attraction is dominated by two characteristic times, depending on the absolute value of the Floquet multipliers. A rapid response happens for *w* and *q*, while a slower response resembling a phugoidal mode [19, 32], with period of about 8*s* characterizes the trends of *u* and *θ*.

**Figure 7:**
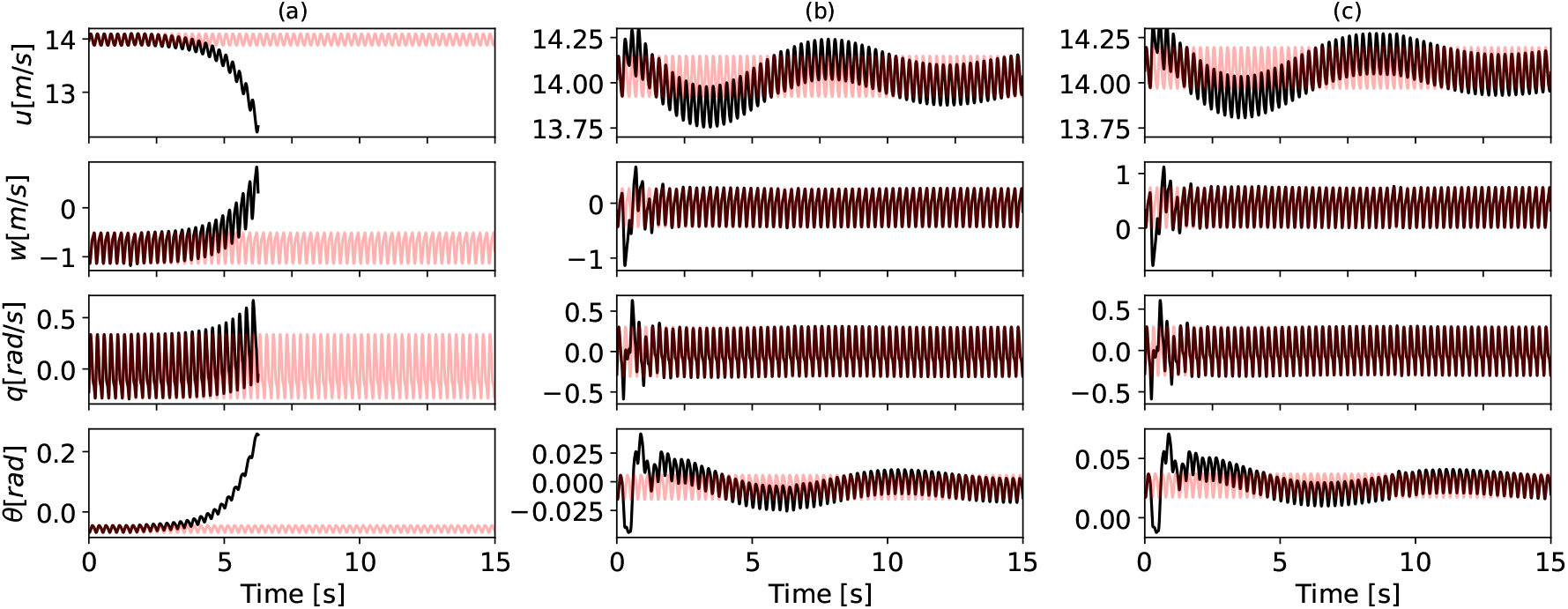
Dynamic response of the three representatives limit cycles, to a Gaussian-like upward gust. **Case (a):** Separation of the perturbed solution along the unstable eigendirection (black) from the periodic orbit. **Case (b)** and **(c):** Passively stable response. After a transient, the perturbed trajectory (black) tends to converge back to the leveled steady flight condition (red).

### iii. Trade-off between CoT and flight stability

Figure 8 further illustrates a trade-off between stability and CoT. Figure 8(a), illustrates the lowest achievable CoT as a function of the forward flight velocity. This is represented at four different values of tail opening *β* = [0, 15, 30, 45]deg. The minimum of the four curves is around 0.085 and corresponds to forward flight velocities of approximately 11*ms*^−1^. The steepness of the curve at increasing velocities monotonically increases with the tail opening. For the same values of *β*, Figure 8(b) reports the Pareto front of the largest Floquet multiplier Λ and CoT. This front captures the optimal solutions for which one of these features could not be more favorable without negatively affecting the other. The transition between stable and unstable flight regimes is highlighted by the vertical purple line.

**Figure 8:**
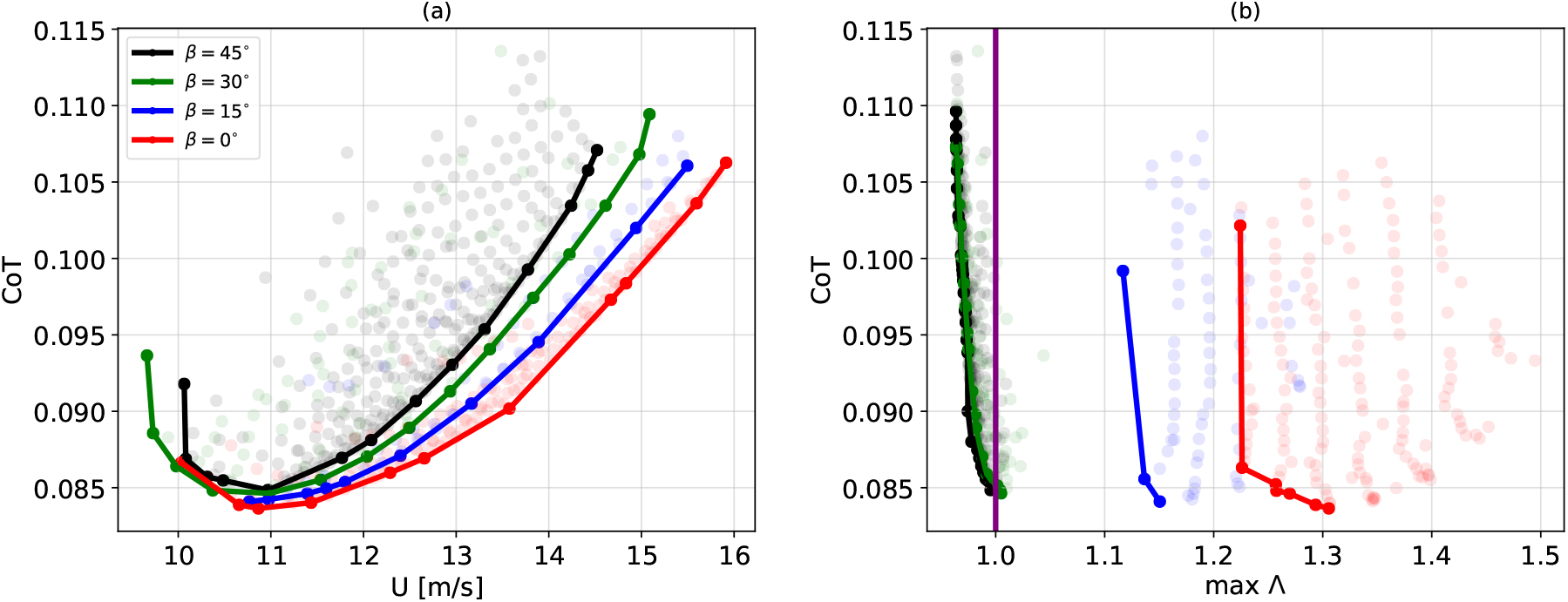
Trade-off between CoT and stability. (a): Lower envelope of the CoT as a function of the forward flight velocity of four evenly-spaced β-planes. We report all the possible solutions, from which the lower envelope is extracted, with transparent points, colored accordingly with the respective tail opening. (b): Pareto front of the CoT as a function of the largest Floquet multiplier of four evenly-spaced β-planes. We report all the possible solutions, from which the Pareto front is extracted, with transparent points, colored accordingly with the respective tail opening. The stability transition is highlighted with the purple vertical line.

## V. Discussion and conclusion

We performed a four-dimensional bifurcation study in the parametric space of flapping gaits. Our numerical analysis highlights the existence of two sets of solutions as a function of the tail opening, with their respective stability properties. Figure 5 shows that steady leveled flight can be achieved for a large set of parameter combinations. Such combinations have to balance the pitching moment generated by the wing and the tail. This condition is mainly driven by the sweep offset of the wing and the angle of tail opening. Both of these parameters indeed modulate the distribution of nose-up and nose-down moment, and thus play a fundamental role in the limit cycle stability. The shoulder amplitude only marginally affects stability, confirming the results reported in [13]. This is due to the fact that it does not have an effect in moving the aerodynamic forces forward or backward — on average — with respect to the center of mass, and thus in altering the pitching moment distribution.

Two profiles of pitching moment that guarantee a passively stable limit cycle have been found. One configuration is similar to those guaranteeing static pitch stability in aircraft and in bird gliding [5, 19], i.e. with the wings generating nose-down moment on average, and the tail generating nose-up moment on average (Figure 6(b)). The second configuration guaranteeing stable limit cycles produces a nose-up moment on average with the wing, and nose-down moment with the tail. These two stable configurations previously discovered for gliding [8], are shown here to also apply to flapping of medium to large-size birds. In [3], Smith stated a biological intuition that birds lost the capacity to rely on passively stable configurations while developing sensory-driven neural circuitries to actively control their flight over the course of evolution. However, this has been recently challenged for gliding flight [5, 7]. It was shown that in gliding regimes, birds can modulate the elbow sweep to achieve passive stability. Here, we extend this promise thesis to flapping regimes, showing that passive stability can also be achieved with appropriate wing kinematics, and tail opening. However opening the tail comes with an important additional energetic cost. A power analysis revealed that this additional energetic cost is due both to overcome the extra drag produced by the tail, but also to the intrinsic efficiency of the adopted wing kinematics leading to the same flight velocity. Figure 6, indeed shows that the extra power required to operate with the open tail conditions, is not only due to drag forces produced by the tail itself, but also to a more costly kinematics adaptation of the gait to level the flight.

The stability of these three conditions has been analyzed under a Gaussian-like gust perturbation. The unstable case shows a quick separation from the limit cycle condition. Interestingly, in the stable solutions, two characteristic times appears: a fast mode of response, affecting the variable *w, q*, and a slow phugoidlike mode that affects *u* and *θ*.

The trade-off between stability and energetic performance is highlighted in Figure 8. The lower envelope of the CoT shows a monotonic increase of curvature with the tail opening. For forward flight velocities of 14*ms*^−1^ the saving in terms of CoT between a furled tail configuration and a full open tail is of about 10%. This is comparable to the energetic advantage drawn from formation flight with respect to solo flight, according to [33]. Indeed, CoT curves with small curvatures are crucial for long range flights, as it allows to modulate the velocity at a lower energetic cost. Figure 8(b) shows the Pareto front of the CoT with respect to the largest Floquet multiplier. The Pareto front of the stable solutions (black and green) is very steep, suggesting that little advantages of stability gain comes with a disproportionate energetic cost. We infer the existence of a close interplay between stability and energetic cost of flapping for medium to large size birds in steady flight. Indeed, whereas the absolute minimum CoT only marginally changes as a function of the tail open-ing, its large variation with respect to the forward velocity does vary as a function of this angle. Put differently, the typical U-shaped curve characterizing the CoT as function of the forward flight velocity is found at each tail opening angle, but its asymptote is smaller for smaller angles.

In this study, we focused on wing and tail contributions to the longitudinal dynamics. To increase the model fidelity, it will be necessary to account other morphological and biological elements that may contribute to stabilize the flight. These would need a substantial adaptation of the equations of motion used in the current work. Moreover, in the current version of our model the wings are assumed to be rigid and no kinematic adaptation is implemented to react to a perturbation. This should be relaxed in a more biocompatible version of the model, that should account for the intrinsic joint compliance due to actuation by muscle-tendon units. This will definitively influence the dynamics of the response to perturbation such as during gust alleviation. Aeroeleasticity along the wingspan should have a similar influence and will be studied in future work [34].

We used a carefully formulated framework to study steady flight stability, i.e. Floquet theory combined with a multiple shooting algorithm. We concluded that in spite of the gain in stability, having a tail-like surface determines an increase of steepness of the CoT with respect to the forward flight velocity, that limits the authority of the bird to modulate the flight speed. This suggests an explanation for the field observation that birds flap with furled tails in long flights [35], i.e. that a loss of dynamic stability might be traded-off in exchange for freedom of modulating velocity at lower energetic expenditure. This might prove to be a crucial factor, for instance, in seasonal migrations, where the time of arrival at foraging, breeding and wintering sites is naturally constrained by environmental factors such as daylight duration, food availability or social reasons. However, our results show that birds still have the authority to select passively stable modes —i.e. with an open tail — that may prevail in certain circumstances, such as flying while sleeping [36].

## A. Appendix

The bird parameters to capture the morphology of the northern bald ibis are reported in Table 2. The data related to the bird morphology are listed in Table 2(a). The parameters governing the wing kinematics after solving Equation 2 are reported in Table 2(b).

**Table 2:** Numerical parameters used in the simulations. Parameters highlighted with an asterisk * are those being varied in the parametric study

| Body |  |
| --- | --- |
| Mass ($m_b$), [kg] | 1.2 |
| Moment of Inertia ($I_{yy}$), [$kg \cdot m^2$] | 0.1 |
| Wing |  |
| Wingbeat frequency $f$, [Hz] | 4 |
| Wingspan ($b$), [m] | 1.35 |
| Mean aerodynamic chord ($\bar{c}$), [m] | 0.15 |
| Arm bone length ($l_a$), [m] | 0.134 |
| Forearm bone length ($l_f$), [m] | 0.162 |
| Hand bone length ($l_h$), [m] | 0.084 |
| Feathers |  |
| Primary feather 1 ($l_{k1}$), [m] | 0.25 |
| Primary feather 2 ($l_{k2}$), [m] | 0.275 |
| Primary feather 3 ($l_{k3}$), [m] | 0.25 |
| Secondary feather 1 ($l_{k4}$), [m] | 0.225 |
| Secondary feather 2 ($l_{k5}$), [m] | 0.2 |
| Secondary feather 3 ($l_{k6}$), [m] | 0.175 |
| Secondary feather 4 ($l_{k7}$), [m] | 0.15 |
| Tail |  |
| Tail chord ($c_0$), [m] | 0.25 |
| Tail opening ($\beta$), [$^\circ$] | $\beta^*$ |

| Joint | $q_0$ [deg] | $A$ [deg] | $\phi$ [deg] |
| --- | --- | --- | --- |
| Shoulder $y$ | 11.5 | 0.8 | -90 |
| Shoulder $x$ | 0 | $A_{s,x}^*$ | 180 |
| Shoulder $z$ | $\psi_{s,z}^*$ | 20 | 90 |
| Elbow $x$ | 30 | 30 | -90 |
| Wrist $y$ | $q_{w,y}^*$ | 30 | -90 |
| Wrist $z$ | -30 | 30 | 90 |

## Acknowledgment

This work was supported by Fédération Wallonie-Bruxelles (FWB) under the Action de recherche concertée (ARC) RevealFlight (grant number **17/22-080, REVEALFLIGHT** – The reverse-engineering of flight: a bottom-up reproduction of bird biomechanics and of self-organization into a flock).

## Code availability

The calculations are performed with the in-house and open source Python package multiflap. The code is publicly available at https://github.com/vortexlab-uclouvain/multiflap.

